# dia-PASEF Proteomics of Tumor and Stroma LMD Enriched from Archived HNSCC Samples

**DOI:** 10.1101/2024.08.09.607341

**Authors:** Aswini Panigrahi, Allison L Hunt, Diego Assis, Matthew Willetts, Bhaskar V Kallakury, Bruce Davidson, Thomas P Conrads, Radoslav Goldman

**Affiliations:** Department of Oncology, Lombardi Comprehensive Cancer Center, Georgetown University, Washington, DC 20057, USA; Women’s Health Integrated Research Center, Women’s Service Line, Inova Health System, Annandale, VA 22003, USA; Bruker Scientific, Billerica, MA 01821, USA; Department of Pathology, Lombardi Comprehensive Cancer Center, Georgetown University, Washington, DC, United States; Department of Otolaryngology-Head and Neck Surgery, Medstar Georgetown University Hospital, Washington, DC, United States; Department of Biochemistry and Molecular & Cellular Biology, Georgetown University, Washington, DC 20057, USA

## Abstract

We employed laser microdissection to selectively harvest tumor cells and stroma from the microenvironment of formalin-fixed, paraffin-embedded head and neck squamous cell carcinoma (HNSCC) tissues. The captured HNSCC tissue fractions were analyzed by quantitative mass spectrometry-based proteomics using a data independent analysis approach. In paired samples, we achieved excellent proteome coverage having quantified 6,668 proteins with a median quantitative coefficient of variation under 10%. We observed significant differences in relevant functional pathways between the spatially resolved tumor and stroma regions. Our results identified extracellular matrix (ECM) as a major component enriched in the stroma, including many cancer associated fibroblast signature proteins in this compartment. We demonstrate the potential for comparative deep proteome analysis from very low starting input in a scalable format that is useful to decipher the alterations in tumor and the stromal microenvironment. Correlating such results with clinical features or disease progression will likely enable identification of novel targets for disease classification and interventions.

## Introduction

Head and neck squamous cell carcinoma (HNSCC) is the sixth most common cancer worldwide accounting for approximately 890,000 new cases and 450,000 deaths (4.6% of cancer deaths) annually (Barsouk et al., 2023; Johnson et al., 2020). HNSCC cells first invade the basement membrane of native epithelium; a large proportion of patients are identified at the time of diagnosis to have lymph node metastasis, which is associated with poor survival (Johnson et al., 2020; Leusink et al., 2018; Sanderson and Ironside, 2002). Overall, the response to available treatments has been moderate. Linking cellular phenotypes to functional proteome states of the HNSCC tumors, including its microenvironment, will add to our understanding of the patho-physiology of the disease and potentially identify drivers of metastasis.

In an earlier study, laser microdissection (LMD) of HNSCC FFPE samples in combination with quantitative mass spectrometry (MS) was introduced by us to characterize normal and tumor lesions (Patel et al., 2008). Recent deep proteomic analysis of tumor and matched normal adjacent tissues, and integrated proteogenomic analysis of MS-based proteomics data with genomics and transcriptomics, have identified molecular subtypes with treatment potential (Huang et al., 2021). However, the proteome coverage of early stage tumors remains insufficient, in part due to the limited availability of surgical tissues for molecular analysis at this early stage of the disease. Likewise, the spatial resolution of the available HNSCC proteomic data and its correlation with clinico-pathological features is very limited. Recent advances in the LMD technology and MS-based techniques allow deep proteome analysis from very low input peptide samples (Demichev et al., 2022; Hunt et al., 2021; Meier et al., 2020; Mitchell et al., 2022; Mund et al., 2022; Truong et al., 2024).

Motivated by these advancements, in this study, we separately harvested tumor and stroma regions from HNSCC FFPE samples using LMD for independent MS-based proteomics analysis using a data independent analysis (DIA) approach. We achieved deep proteome coverage, identified significant differences in functional pathways of tumor cells and the stromal microenvironment, and demonstrated the potential for comparative deep proteome analysis in a scalable format.

## Results and Discussion

### dia-PASEF LC-MS analysis of tumor and stroma region

Paired tumor and stroma regions were enriched by LMD from five HNSCC patient FFPE tissue thin sections that were independently analyzed by DIA MS (Figure 1A, B). The samples were processed, digested with trypsin, and analyzed in triplicate (500 ng peptide digest per injection) on a timsTOF HT MS (40 min run time equating to a ∼2 h MS analytical time per sample). Excellent chromatographic reproducibility was observed over the entire sample batch run (Figure 1C, D). Taken together, a total of 8,759 protein groups were identified from 98,005 peptide matches among which 8,498 proteins were from LMD enriched tumor epithelium and 8,396 proteins from the stromal microenvironment, with an overlap of 94% between the two histologically distinct tissue regions. Of these proteins, 7,825 were identified with 2 or more peptide matches in at least one of the samples (Supplementary Table 1A). We observed similar protein coverage (range 7,543 to 7,709) in each sample (Figure 1E). Across the patient samples 6,668 proteins were co-quantified (Supplementary Table 1B), with the median CVs consistently under 10% (Figure 1F). This analysis provides an in-depth snapshot of the proteome in the enriched stromal and tumor regions with significant improvements in both the coverage and quantification of HNSCC patient tissues.

**Figure 1.**
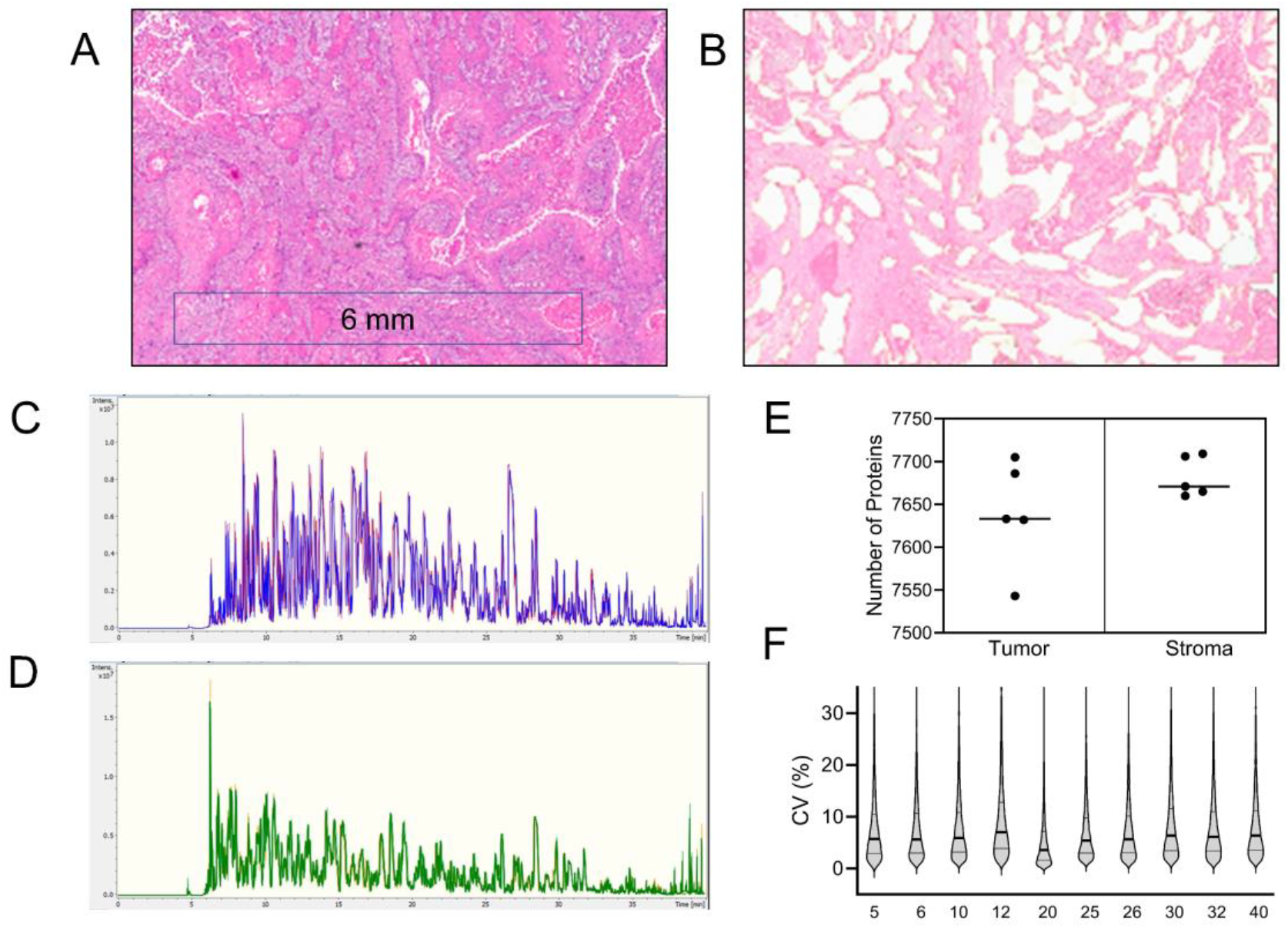
Representative (A) H&E stained FFPE tissue section, and (B) image after LMD capture of the tumor regions. Base peak chromatograms showing the overlay of triplicate runs of an enriched (C) tumor and (D) paired stroma sample. (E) Graphical representation of number of proteins identified with 2 or more peptide matches in the enriched tumor and stroma samples. The median is indicated by a line. (F) Schematic representation of CV value spread of quantitation for each sample, median (bold line) and upper and lower quartile (thin line) are indicated.

### Differential proteome in tumor and stroma

The samples clustered as two separate groups (stroma and tumor) in hierarchical analysis (Figure 2A). A differential analysis identified 2,655 significantly altered proteins between tumor epithelium and stroma (Figure 2B). A gene ontology molecular functional pathway enrichment analysis of these proteins (Supplementary Table 2) identified several functional groups relevant to the stromal and tumor regions (Figure 2C, D). Notably extracellular matrix (ECM) structural constituent, glycosaminoglycan binding, heparin binding, collagen binding, and calcium ion binding are more abundant in stroma; and nucleic acid catalytic activity, DNA binding, ATP hydrolysis, helicase and ribosome constituents are more abundant in the tumor epithelium proteome. This suggests nearly exclusive expression pattern in the tumor or stromal regions, and the enrichment is efficient and allows improved quantitative analysis of stromal features compared to the bulk tissue analysis.

**Figure 2.**
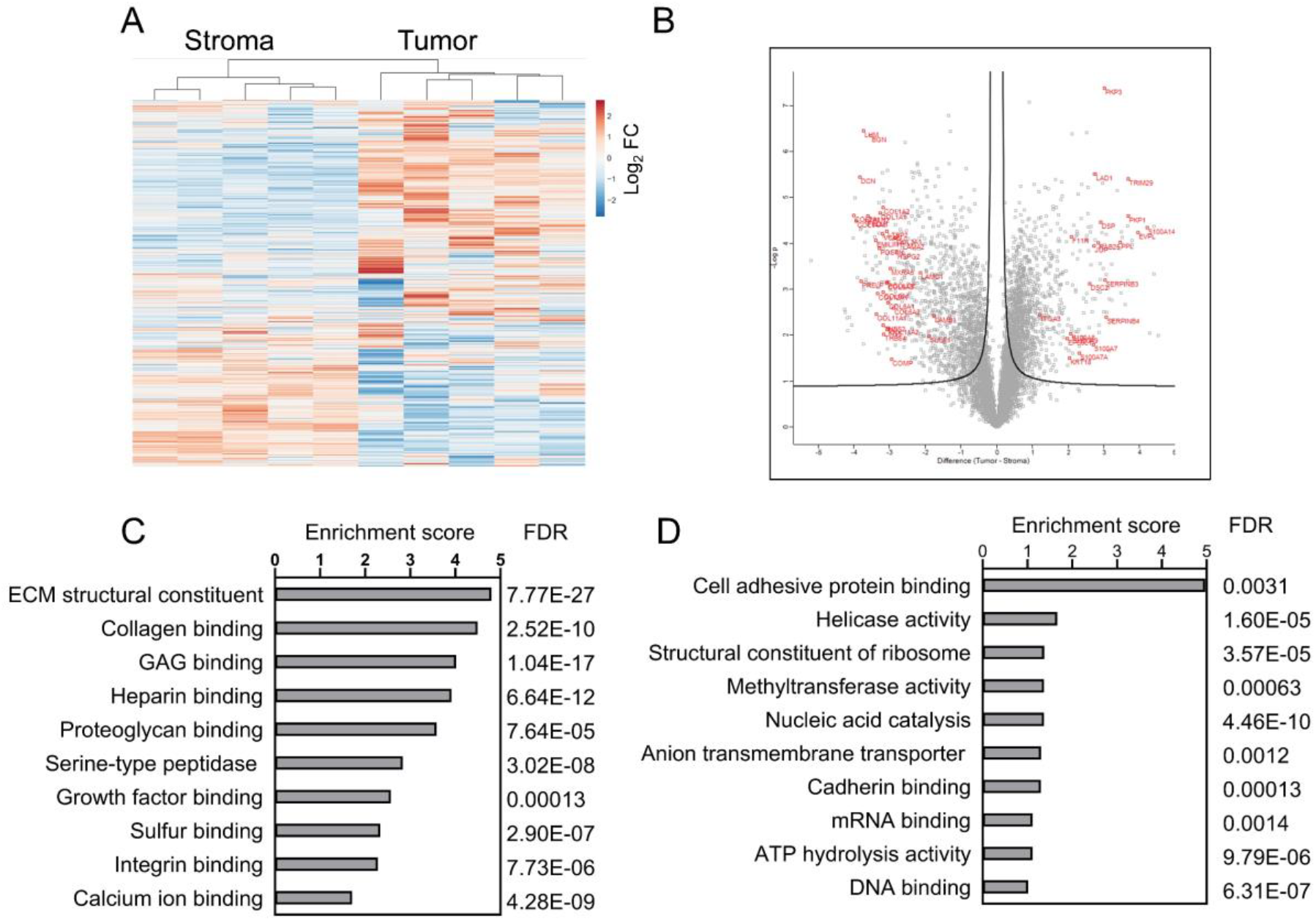
Functional association of upregulated proteins. (A) Unsupervised hierarchical cluster analysis of stroma and tumor region proteins based on abundance, and (B) Volcano plot of protein abundance. In volcano plot the box dot plots of significantly changed proteins between the regions are indicated, and selected candidates are in red. Selected enriched functional pathways (String analysis) in stroma (B) and tumor (C) are shown with enrichment score and false discovery rate (FDR).

### Extracellular matrix organization

ECM remodeling is clearly enriched in the stromal microenvironment. Several components of ECM structural constituent, including BGN, COL11A1, COL11A2, COL12A1, COL14A1, COL1A1, COL1A2, COL2A1, COL3A1, COL6A1, COL6A2, COL6A3, COL8A1, COMP, DCN, EMILIN1, FBLN1, FBLN2, FN1, HSPG2, LUM, MXRA5, PRELP, and VCAN, were among the top 100 most abundant proteins in stroma. Some of the proteins with higher abundance in stroma (e.g. COL11A1, COMP, FN1, POSTN, SULF1 and THBS2) are signature markers of COL11A1-expressing cancer associated fibroblasts (CAFs) (Zhu et al., 2021) detectable in several cancers including HNSCC (Yang et al., 2022). This type of CAF was described as a common driver of aggressive cancer behavior (Zhu et al., 2021), and we present a functional and physical protein association map of these CAF associated proteins (that are more abundant in stroma) in Fig 3A. The results are in agreement that CAFs are enriched in the stromal microenvironment that serve as the primary source of ECM (Li et al., 2024).

**Figure 3.**
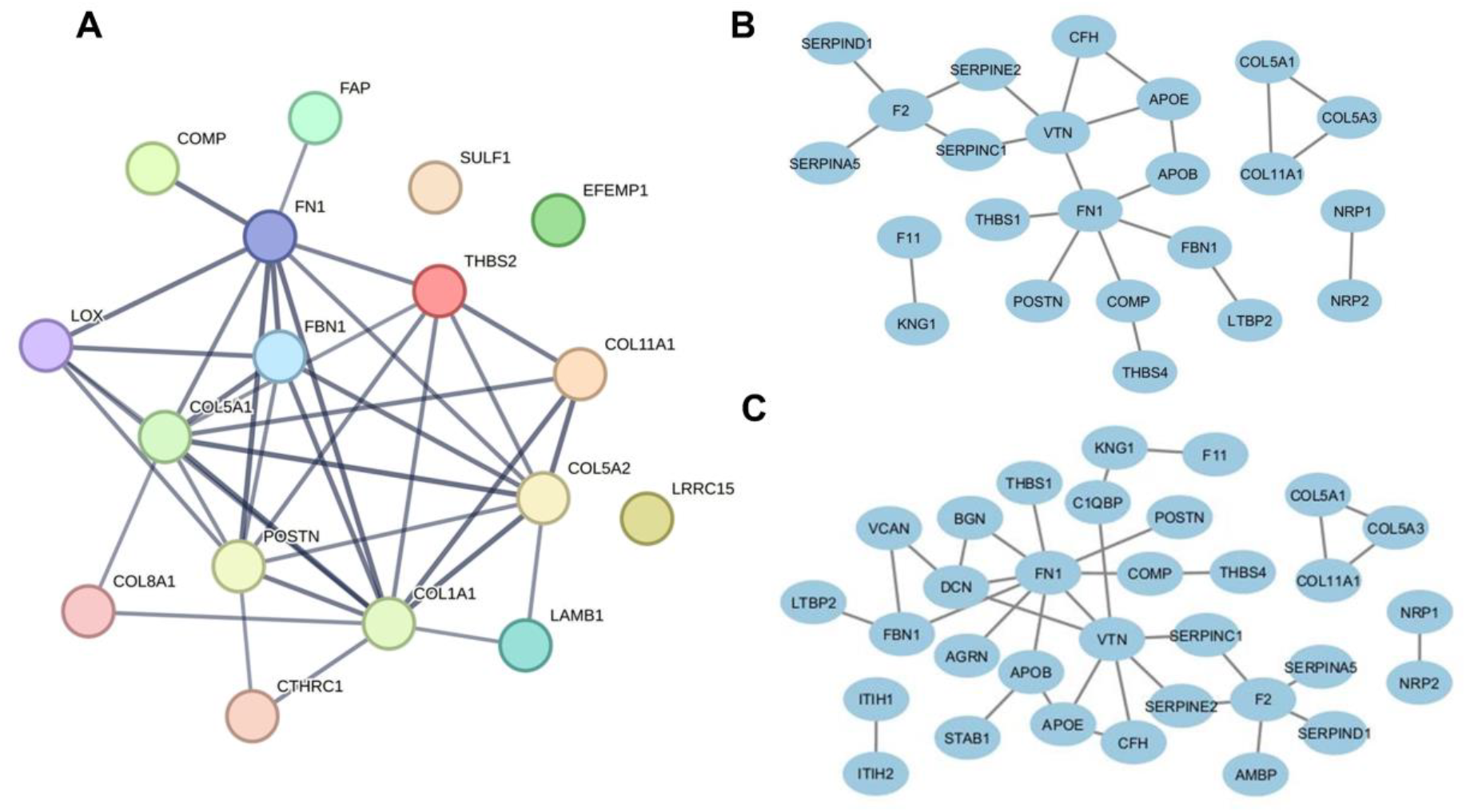
High confidence STRING network (A) functional and physical association of upregulated CAF markers in stroma, physical association of upregulated stromal (B) heparin binding and (C) GAG binding proteins.

We also identified a set of heparin binding proteins to be significantly enriched in the stromal microenvironment, including COL11A1, COMP, FN1, LTBP2, PCOLCE, POSTN, THBS4, THBS2, and PRELP. A high confidence physical interaction map of stromal enriched heparin binding proteins is shown in Fig 3B. We also present an interaction map of glycosaminoglycan (GAG) binding proteins (Fig 3C), and some of these proteins overlap with heparin binders. KEGG pathway enrichment analysis of all ECM component proteins in our dataset showed association with ECM-receptor interaction, protein digestion and absorption, small cell lung cancer, focal adhesion, relaxin signaling, PI3K-Akt signaling, IL-17 signaling, TGF-beta signaling, and proteoglycans in cancer (Table 1). Some of the ECM-receptor proteins (e.g. laminin subunits LAMA2, LAMB1, and LAMC1) that are more abundant in stroma may bind to cells via high affinity receptors and mediate cell attachment, migration, and organization.

**Table 1.**
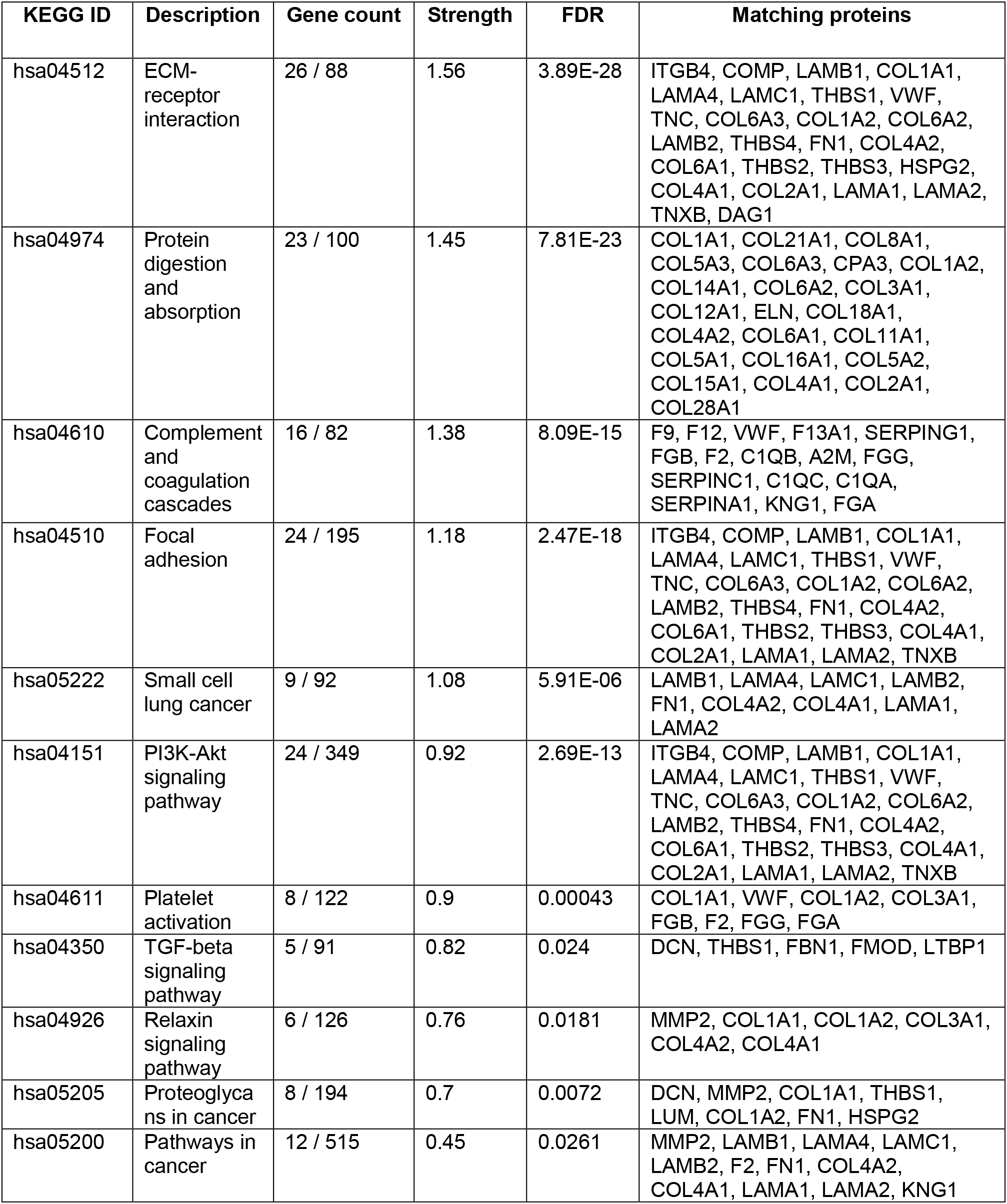
KEGG pathway enrichment of ECM proteins that changed significantly between stromal and tumor region.

| KEGG ID | Description | Gene count | Strength | FDR | Matching proteins |
| --- | --- | --- | --- | --- | --- |
| hsa04512 | ECM-receptor interaction | 26 / 88 | 1.56 | 3.89E-28 | ITGB4, COMP, LAMB1, COL1A1, LAMA4, LAMC1, THBS1, VWF, TNC, COL6A3, COL1A2, COL6A2, LAMB2, THBS4, FN1, COL4A2, COL6A1, THBS2, THBS3, HSPG2, COL4A1, COL2A1, LAMA1, LAMA2, TNXB, DAG1 |
| hsa04974 | Protein digestion and absorption | 23 / 100 | 1.45 | 7.81E-23 | COL1A1, COL21A1, COL8A1, COL5A3, COL6A3, CPA3, COL1A2, COL14A1, COL6A2, COL3A1, COL12A1, ELN, COL18A1, COL4A2, COL6A1, COL11A1, COL5A1, COL16A1, COL5A2, COL15A1, COL4A1, COL2A1, COL28A1 |
| hsa04610 | Complement and coagulation cascades | 16 / 82 | 1.38 | 8.09E-15 | F9, F12, VWF, F13A1, SERPING1, FGB, F2, C1QB, A2M, FGG, SERPINC1, C1QC, C1QA, SERPINA1, KNG1, FGA |
| hsa04510 | Focal adhesion | 24 / 195 | 1.18 | 2.47E-18 | ITGB4, COMP, LAMB1, COL1A1, LAMA4, LAMC1, THBS1, VWF, TNC, COL6A3, COL1A2, COL6A2, LAMB2, THBS4, FN1, COL4A2, COL6A1, THBS2, THBS3, COL4A1, COL2A1, LAMA1, LAMA2, TNXB |
| hsa05222 | Small cell lung cancer | 9 / 92 | 1.08 | 5.91E-06 | LAMB1, LAMA4, LAMC1, LAMB2, FN1, COL4A2, COL4A1, LAMA1, LAMA2 |
| hsa04151 | PI3K-Akt signaling pathway | 24 / 349 | 0.92 | 2.69E-13 | ITGB4, COMP, LAMB1, COL1A1, LAMA4, LAMC1, THBS1, VWF, TNC, COL6A3, COL1A2, COL6A2, LAMB2, THBS4, FN1, COL4A2, COL6A1, THBS2, THBS3, COL4A1, COL2A1, LAMA1, LAMA2, TNXB |
| hsa04611 | Platelet activation | 8 / 122 | 0.9 | 0.00043 | COL1A1, VWF, COL1A2, COL3A1, FGB, F2, FGG, FGA |
| hsa04350 | TGF-beta signaling pathway | 5 / 91 | 0.82 | 0.024 | DCN, THBS1, FBN1, FMOD, LTBP1 |
| hsa04926 | Relaxin signaling pathway | 6 / 126 | 0.76 | 0.0181 | MMP2, COL1A1, COL1A2, COL3A1, COL4A2, COL4A1 |
| hsa05205 | Proteoglycans in cancer | 8 / 194 | 0.7 | 0.0072 | DCN, MMP2, COL1A1, THBS1, LUM, COL1A2, FN1, HSPG2 |
| hsa05200 | Pathways in cancer | 12 / 515 | 0.45 | 0.0261 | MMP2, LAMB1, LAMA4, LAMC1, LAMB2, F2, FN1, COL4A2, COL4A1, LAMA1, LAMA2, KNG1 |

The ECM is dynamic, provides signals regulating the tumor cells, and impacts their migration and metastatic potential (Kai et al., 2019). Characterization of the stromal microenvironment in HNSCC tissues is valuable (Li et al., 2024; Mukherjee et al., 2023; Principe et al., 2018) and will likely allow connection of disease pathology to outcomes using the pathology guided LMD enrichment coupled with the high sensitivity of LC-MS/MS using minute amounts of input starting material.

### Proteins enriched in the tumor fractions

In the top 100 tumor enriched proteins we identified candidates with functional and physical association with cell-cell adhesion, including DSC2, DSP, JUP, KRT18, PKP3, and TRIM29. We also identified cadherin binding (EPS8L1, EVPL, F11R, JUP, KRT18, LAD1, PKP1, PKP3, PPL, TRIM29) and calcium-dependent protein binding (S100A14, S100A7, S100A7A, S100A8, S100A9) proteins.

Some of the major changes in the tumor epithelium proteome were related to nucleic acid binding and processing (Supplementary Table 2), and they are associated with DNA replication, mismatch repair, and RNA processing. These data, along with upregulated ribosome biogenesis in tumor, may reflect a higher cell proliferation rate compared to stroma. By keyword search we identified proteins in this dataset associated with cell invasion; including integrin ITGA3 enriched in the tumor epithelium, which may provide a docking site for FAP at invadopodia plasma membranes in a collagen-dependent manner, and hence may participate in the adhesion, formation of invadopodia, and matrix degradation processes, promoting cell invasion. Some of the highly enriched proteins in the tumor epithelium, including RAB25, may promote the invasive migration of cells, and SERPINB3 and SERPINB4 may modulate the host immune response against tumor cells.

Overall, we demonstrated excellent proteome coverage and relative quantitation of HNSCC tumor and its microenvironment using LMD in combination with a highly sensitive and quantitative dia-PASEF LC-MS/MS workflow. Though our sample set was limited, we document the expected changes at spatial level occurring in stroma adjacent to the tumor. We used 1.5 μg of isolated tryptic peptides for triplicate analysis per tissue specific region, which allows an efficient quantitative protein analysis of the LMD enriched FFPE tumor and stromal regions. The methodology allows comparative deep proteome analysis in a scalable format and allows connection of the enriched proteomes to pathologic features of the tumor tissues. Even though the protein yield from the stromal regions is lower, the sensitivity of the LC-MS/MS workflow allows reproducible proteome analysis with coverage comparable to the enriched tumor regions. Our future focus is on expansion of this study in a large sample cohort using this workflow that would allow us to identify proteins relevant to disease progression and survival.

## Materials & Methods

### Experimental Design and Statistical Rationale

In this study, proteomic profiling of five patient samples was conducted. From each sample the tumor and stroma enriched region was captured by LMD. LC-MS analysis of these paired samples was performed in triplicates to demonstrate the technical reproducibility, and statistical analysis was performed to determine the CV for each sample analysis. The quantitation values between the two paired sample groups, i.e. tumor enriched (n=5) and stroma enriched (n=5) were compared, and Student’s T-test was performed to capture statistically significant relative protein differences.

### Patient specimens

Study participants were enrolled between 1995 and 2020 at the Department of Otolaryngology-Head and Neck Surgery, MedStar Georgetown University Hospital, under a protocol approved by the MedStar Health Research Institute-Georgetown University Oncology Institutional Review Board. Thin sectioned formalin-fixed paraffin-embedded (FFPE) HNSCC tumor specimens mounted on polyethylene naphthalate membrane slides were prepared by Histology & Tissue Shared Resource at Georgetown University Medical Center. The samples were stained with hematoxylin and eosin (H&E), imaged by microscopy, and reviewed by board-certified pathologist to annotate the tumor and stroma areas.

### Laser capture microdissection

LMD of tumor and stromal regions was performed using methods recently described and imaged before after the capture (Hunt et al., 2021). The samples were prepared for mass spectrometry analysis by trypsinization and the peptides were desalted using C18 cartridges.

### LC-MS analysis

A timsTOF HT mass spectrometer connected to a nanoElute 2 LC system via the CaptiveSpray 2 source (Bruker, Billerica, MA) was used for dia-PASEF LC-MS/MS analysis. Each sample was analyzed in triplicate (except one sample in duplicate, that was used for initial method optimization and availability was limited). The peptides were separated by a C18 IonOpticks column (particle size 1.6 μm, 75 uM μm ID, 25 cm length) at flow rate of 0.25 μL/min; solvent A (0.1% formic acid in water), solvent B (0.1% formic acid in Acetonitrile), gradient of 0 to 28 min 5-23 % B, 28 to 32 min to 30% B, 32 to 36 min to 90% B, and hold at 90% B to 40 min. For dia-PASEF analysis the window scheme was calculated using the py_diAID tool (https://github.com/MannLabs/pydiaid). The capillary voltage was set at 1600 V, stepping collision energy at 32, 40, 50 eV, dia-PASEF scan range 100-1700 *m/z* in positive mode, and IMS service ramp time of 100 ms.

### Data analysis

Acquired mass spectrometry data was searched against Uniprot-Human-reviewed database (20,383 protein entries) using the directDIA+ workflow (Spectronaut 18 software) for protein identification and quantification using BGS default settings. Unsupervised hierarchical clustering analyses was done using Clustvis (v 1.2.0) in R package (v 3.6.2) (Metsalu and Vilo, 2015). Perseus software (v 2.0.11) was used to compare the relative abundance of two sample groups, perform Student’s T-test and Volcano plot analysis (Tyanova et al., 2016). Molecular functional pathway enrichment analysis was performed using String DB (v 12.0) for proteins showing statistically significant variation between the two groups. Cytoscape (v 3.10.2) was used to generate schematic output (Shannon et al., 2003) from STRING network analysis. GraphPad Prism Software (v 10.2.3) was used for data visualization.

## Author Contributions

AP, TPC, RG writing, review and editing; TPC, RG supervision and resources; RG funding acquisition; AP, ALH, TPC, RG conceptualization; AP, ALH, DA, MW, BVK, BD methodology, investigation and formal analysis; AP, ALH, DA data curation and visualization.

## Conflict of interest

AP, ALH, BVK, BD, TPC, RG declare that they have no known competing financial interests. DA, MW are employees of Bruker Scientific.

## Acknowledgements

We thank Julius Benicky and Pritha Mukherjee, Georgetown University for helpful suggestions. This work was supported in part by a National Institutes of Health grant R01CA238455 to RG. Further support was provided by Georgetown University Lombardi Comprehensive Cancer Center Support Grant from National Cancer Institute 2P30CA051008. The content is solely the responsibility of the authors and does not necessarily represent the official views of the National Institutes of Health.

## Data Availability

The mass spectrometry data have been deposited to the jPOST repository (Okuda et al., 2017). The accession numbers are PXD054650 for ProteomeXchange and JPST003262 for jPOST (https://repository.jpostdb.org/preview/140548518366b29246602a8 Access key: 1101)

## Notes

### Competing Interest Statement

Authors AP, ALH, BVK, BD, TPC, RG declare that they have no known competing financial interests. DA, MW are employees of Bruker Scientific.

